# Exome capture of Antarctic krill (*Euphausia superba*) for cost effective population genetics of historical collections

**DOI:** 10.1101/2023.07.24.550387

**Authors:** Oliver White, Geraint Tarling, Lauren Hughes, Sarah Walkington, Matt Clark

## Abstract

Antarctic Krill (*Euphausia superba*) is a pivotal keystone species in the Southern Ocean ecosystem, with immense ecological and commercial significance. However, its vulnerability to climate change necessitates urgent investigation of its population genetics and adaptive responses. Historical spirit collections of Antarctic krill from the early 20th century represent an ideal opportunity for genomic research, to investigate how krill have changed over time and been impacted by predation, fishing and climate change. In this study, we assessed the utility of shotgun sequencing and exome capture for genomic analyses with historical spirit collections of Antarctic krill. Because the krill genome is very large (48Gb) two full-length transcriptomes were generated and used to identify putative targets for targeted resequencing. Skim genome sequencing allowed sample and library quality control. By comparing genome to exome resequencing of the same libraries we calculate enrichment and variant calling metrics. Full-length mitochondrial and nuclear ribosomal sequences were successfully assembled from genomic data demonstrating that endogenous DNA sequences could be assembled from historical collections. We find that exome capture provided enrichment of on-target sequence data, with increased depth and higher variant quality for targeted loci. Our findings demonstrate the feasibility of extracting genomic information from historical krill samples, despite the challenges of fragmented DNA and huge genome size unlocking such collections to provide valuable insights into past and present krill diversity, resilience, and adaptability to climate change. This approach unlocks the potential for broader genomic studies in similar samples, and for enhancing conservation efforts and fisheries management in the Southern Ocean ecosystem.

## Introduction

Antarctic Krill (*Euphausia superba*) is a keystone species in the Southern Ocean ecosystem. Antarctic Krill (hereafter krill) is also the most successful wild living animal species on the planet in terms of population biomass (300-500 million tonnes; Atkinson et al., 2009), with a circumpolar distribution covering the Southern Ocean ecosystem, and krill swarms forming some of the largest aggregations of animal life. Krill’s significance extends far beyond its sheer abundance. It plays a pivotal role in the food web of the Southern Ocean, as a consumer of phytoplankton and prey for charismatic megafauna including penguins, seals and whales. The Southern Ocean is one of the largest carbon sinks in the world and krill plays a key role in the carbon cycle, removing up to 40 million tonnes annually (Cavan et al., 2019). In addition, it has become increasingly commercially important, supporting the krill trawler industry’s catch of >$200M annually for fish oil dietary supplements and aquaculture feed (Tou et al., 2007).

The polar regions are thought to be most at risk from climatic warming, with increases of more than double the global average (Meredith et al., 2019). Rapid warming is likely to have profound implications for species distributions and ecosystem functioning, especially polar species which have low tolerance to fluctuating temperatures. Critically, krill is a stenothermic species, adapted to a narrow temperature range between -2 and 5°C (McBride et al., 2021), making it particularly sensitive to climate change. Over the past 90 years, the range of Antarctic krill has contracted southward (Atkinson et al., 2019). In addition, models suggest that a temporal shift in seasonal timings of habitat quality may cause disjunctions between krill’s biological timings and the future environment (Veytia et al., 2020).

Understanding the population genetics of krill, and how this species will respond to future climate change is crucial for both ecosystem functioning and fisheries management. For example, there may be distinct sub-populations of krill, with varying susceptibility or adaptation to rapidly changing ecological conditions. An understanding of population structure would aid stock management through rotating fishing quotas across populations, allowing genetic diversity and resilience to be maintained, and avoiding overfishing of subpopulations. For example, fisheries may be targeting populations of krill living away from ice in warmer waters that are more easily fished. But these populations could be most adaptable to climate change. However, to date, we do not fully understand krill population structures, to what degree sub-populations are inter-connected and able to replenish spatial regions adversely affected by, for instance, climatic anomalies such as Southern Annular Modes and El Nino/La Nina effects. Without this understanding, we cannot fully determine how they will tolerate, and how they have responded to, fishing, climate change and other human impacts.

Previous population genetics studies of krill have largely relied upon a small selection of markers from allozyme variation (Fevolden & Schneppenheim, 1989) and mitochondrial DNA (Goodall-Copestake et al., 2010; Zane et al., 1998), but these only consider single mitochondrial genes and were not informative of variation at the genomic level. A reduced representation approach (RAD-seq) has been employed to investigate population structure in Krill (Meyer et al., 2015), but this study was hampered because most genetic data were from multicopy genomic regions. More recently, a chromosome level assembly has been generated for krill (Shao et al., 2023), and resequencing of 75 individuals from multiple populations revealed limited population structure but identified 387 sites associated with environmental variables.

There are limited molecular markers available for krill, likely due to the lack of genomic resources available until recently (Shao et al., 2023). Genome resequencing, whilst useful for population genetics, is likely to be cost prohibitive for large populations studies due to the size of the genome (48Gbp) and the amount of sequence data required to call variants confidently. For example, Shao et al (2023) generated 4TB of short read data for their resequencing of 75 individuals which would not be cost effective if scaled to much larger sampling. Another more cost-effective options would be exome capture, to target the portion of the genome that codes for proteins and is an excellent source of single copy sequences for genetic marker discovery.

Natural History collections offer an invaluable resource to understand how krill will respond to climate change. Historical collections represent time stamps of diversity at the time of collection and offer the opportunity to investigate population change and adaptation over geographical and historical gradients. For example, the Natural History Museum London is home to ca. 20,000 krill accessions, with trawl samples suitable for population study spanning a 130-year time frame of the Anthropocene. The collections are also home to samples of great historical importance, including those collected during Scott’s Discovery expedition in 1908. These represent potential flagship collections to investigate the impact of climate change in a keystone species in the Southern Ocean ecosystem throughout the Anthropocene.

This study investigates the relative utility of shotgun sequencing and exome capture for population genetic analyses with historical spirit collections of Antarctic krill. Target exome sequences are identified using two full-length transcriptomes for krill, generated using PacBio IsoSeq data from recently collected samples. The utility of shotgun sequencing and exome capture is then compared for historical krill spirit collections based on target coverage and variant calling. We hypothesise (1) shotgun sequencing will allow the assembly of mitochondrial genome sequences which are present in high copy number, whilst sequencing depth for nuclear single copy genes will be too low for sequence assembly or population genetics analysis and (2) targeted sequencing will increase the likelihood of nuclear gene assembly with sufficient depth for population genetic analyses.

## Methods

### PacBio IsoSeq sequencing

Samples were collected using an EV65 Net1 and immediately flash frozen within liquid Nitrogen and stored at -80°C. Two samples, one male (29659_1) and one female (29659_4) were shipped to NERC Environmental Omics Facility (Sheffield, UK). Samples were dissected for muscle tissue before isolating RNA. RNA quality and integrity was checked before sequencing. High quality transcripts were generated for each sample using the SMRTlink pipeline.

### Target selection and bait design

The high-quality transcripts generated from PacBio IsoSeq data were used to identify putative target sequences. For high-quality transcripts recovered from each sample, the following analyses were performed: (1) BUSCO search against core Eukaryotic genes (eukaryota_odb10; 10/09/2020 using the transcriptome mode), (2) blastn (Camacho et al., 2009) search against annotated transcript sequences downloaded from KrillDB^2^ (Urso et al., 2022) and (3) Blobtools (Laetsch et al., 2017) analysis to define transcript taxonomy based on a blastn search against the NCBI nucleotide database (nt, downloaded June 2022) and the taxrule “bestsumorder”. Transcripts were prioritised for targeted sequencing if they had a (1) BUSCO hit to a core eukaryotic gene, (2) blast hit description from KrillDB^2^ suggesting a role in environmental responses (i.e., heat, cold, temperature sensitive) or core homeobox genes and finally (3) a sequence identified as being from *Euphuasia superba* based on the blobtools output. Putative targets from each sample were combined and redundant sequences were identified and removed using cd-hit-est with a similarity threshold of 0.8 (Fu et al., 2012). Note we also avoided targeting sequences from the mitochondrial genome or genes with putative repetitive elements (e.g., microsatellites) as these are expected to occur at a higher frequency and will bias the sequencing of these targets.

The putative target sequences were shared with Arbor biosciences and tested for bait design suitability. Specifically, targets were softmasked for simple and low complexity repeats and baits were designed using 80 nucleotide (nt) probes and 4× tiling (i.e., one probe every ∼20nt). A blastn search of baits against the *E. superba* reference genome (CNGB: CNP0001930) and mitochondrial genome (NCBI: NC_040987.1) was performed to identify baits that matched overrepresented sequences (e.g., baits with multiple blast hits in the nuclear genome) or mitochondrial sequences which could reduce recovery of target loci due to a higher concentration of mitochondrial sequences in the starting material and higher sequence recovery.

### Shotgun and exome capture sequencing

A total of 27 samples were collected from the NHM spirit collection with the aim of sampling different station numbers from the early historical samples collected in the 1920s and 30s (Table 1; Figure 1). DNA was isolated for a using a modified method from Ruane and Austin (2017) with double quantities of binding buffer/EtOH. DNA samples were shipped to Daicel Arbor BioSciences (Ann Arbor, United States) where total DNA was quantified via a spectrofluorimetric assay and visualised using the TapeStation 4200 (Agilent) platform with a High Sensitivity D1000 tape. Samples containing high molecular weight DNA or no visible DNA were sonicated to generate an average insert size of approximately 300nt before taking up to 80% of the available mass up to 5ng to a single-stranded library preparation protocol that produces dual-indexed Illumina-compatible libraries. For the shotgun sequence data, 20 libraries (Table 1) were pooled in equimolar ratios and sequenced on a partial NovaSeq 6000 S4 PE150 lane, targeting approximately 1M read pairs per sample. Capture pools were prepared from up to 250ng of 6 libraries per reaction for the historical samples. Each capture pool was dried down to 7 uL by vacuum centrifugation. Captures were performed following the myBaits v5.02 protocol using myBaits custom design with an overnight hybridization and washes at 62°C. The captures were pooled in approximately equilmolar ratios. For the exome capture sequence data, 20 samples (Table 1) were sequenced on the Illumina NovaSeq 6000 platform on a partial S4 PE150 lane to approximately 2M read pairs per library.

**Table 1.** Historical samples from the NHM spirit collection used for shotgun and targeted sequencing with information for collection station number, source (Discovery/William Scoresby), year, net type, depth (start-end depth), location and whether or not the sample was used for sequencing.

| N | Station | Collection | Year | Net | Depth (m) | Location | Latitude | Longitude | Sequenced |
| --- | --- | --- | --- | --- | --- | --- | --- | --- | --- |
| 1 | 1311 | Discovery | 1934 | N100H | 5-0 | Bellingshausen sea | -67.313 | -82.900 | Y |
| 2 | 539 | Discovery | 1930 | n70B | 137-0 | South Shetland Islands | -61.800 | -54.858 | Y |
| 3 | 2365 | Discovery | 1938 | TYFB | 350-200 | South Atlantic | -53.390 | 4.842 | Y |
| 4 | 125 | Discovery | 1926 | N100H | 70 | South Georgia | -53.475 | -36.342 | Y |
| 5 | 9968 | Discovery | 1979 | RMT8M-3 | 980-1990 | NA | NA | NA | Y |
| 6 | 2139 | Discovery | 1937 | N100H | 5-0 | Prydz bay | -56.580 | 95.057 | Y |
| 7 | 539 | WS | 1931 | N100B | 100-0 | South Georgia | -57.692 | -23.200 | Y |
| 8 | 1404 | Discovery | 1934 | N100B | 117-0 | South Georgia | -53.997 | -39.465 | Y |
| 9 | 538 | WS | 1931 | N100B | 97-0 | South Georgia | -57.058 | -24.533 | Y |
| 10 | 1405 | Discovery | 1934 | N100B | 110-0 | South Georgia | -54.002 | -40.123 | Y |
| 11 | 542 | WS | 1931 | N100B | 77-0 | South Georgia | -58.650 | -18.217 | Y |
| 12 | 1297 | Discovery | 1934 | N100H | 5-0 | Amunden sea | -70.420 | -129.262 | Y |
| 13 | 42 | Discovery | 1926 | N7-T | 120-204 | NA | NA | NA | Y |
| 14 | 622 | Discovery | 1931 | N100-B | 155-0 | South Georgia | -59.092 | -36.417 | Y |
| 15 | 643 | Discovery | 1931 | N100B | 93-0 | South Shetland Islands | -61.742 | -56.117 | Y |
| 16 | NA | Discovery | 1926 | HN | 0-1 | NA | NA | NA | Y |
| 17 | 568 | WS | 1931 | N100B | 106-0 | NA | NA | NA | Y |
| 18 | 53 | WS | 1927 | N100H | 0-5 | NA | NA | NA | N |
| 19 | 39 | WS | 1926 | N100H | 87 | South Georgia | -54.133 | -35.717 | N |
| 20 | 30 | WS | 1926 | 100H | 0-5 | South Georgia | -53.571 | -38.604 | N |
| 21 | 36 | WS | 1926 | N100H | 77 | South Georgia | -55.338 | -34.775 | Y |
| 22 | 542 | WS | 1927 | N100H | 70 | South Georgia | -58.650 | -18.217 | N |
| 23 | 43 | WS | 1927 | N100H | 70 | South Georgia | -54.900 | -36.833 | N |
| 24 | 1309 | Discovery | 1934 | N100B | 106-0 | Bellingshausen sea | -67.765 | -89.392 | Y |
| 25 | 53H | WS | 1927 | N100H | 0-5 | NA | NA | NA | N |
| 26 | 53O | WS | 1927 | N100H | 0-5 | NA | NA | NA | N |
| 27 | 658 | Discovery | 1931 | N100B | 120-0 | South Georgia | -53.638 | -40.454 | Y |

**Figure 1.**
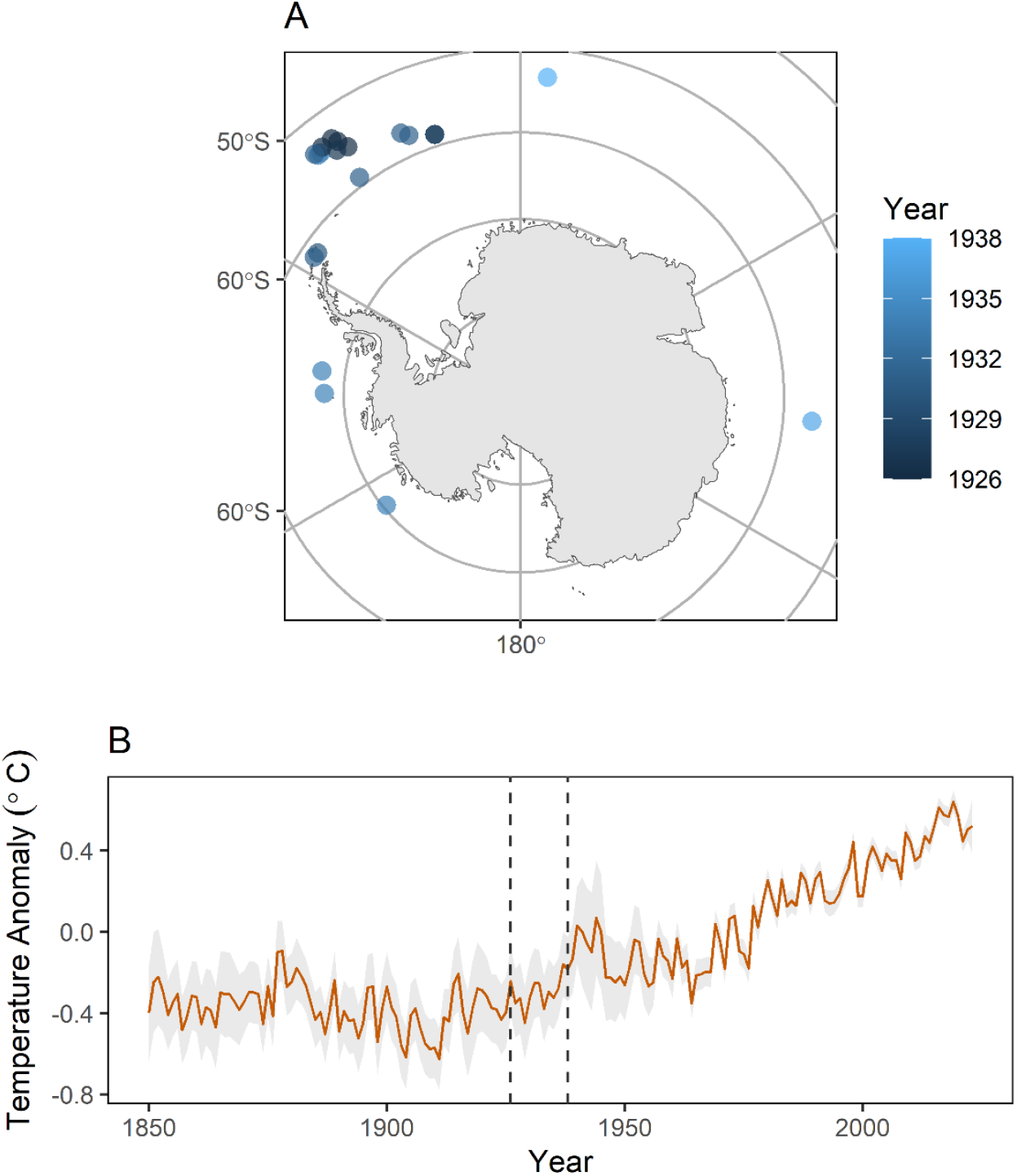
(A) Location of krill (*Euphausia superba*) sampling locations with points coloured by collections year (B) and the Southern Ocean surface temperature relative to 1961-1990 mean, annotated with the minimum and maximum dates collection years of samples used in this study. The sea surface temperature data was downloaded from the Met Office Hadley Centre observations datasets (accessed April 2023).

### Bioinformatics analyses

Raw sequence data generated from shotgun and exome capture were quality filtered using Trimmomatic (Bolger et al., 2014) with the following parameters: ILLUMINACLIP TruSeq3-PE-2.fa 2:30:10, LEADING 3, TRAILING 3, SLIDINGWINDOW 4:15 and MINLEN 36. To evaluate the utility of shotgun data for the assembly multi-copy sequences, GetOrganelle (Jin et al., 2020) was used to assemble mitochondrial genomes and ribosomal genes. Custom GetOrganelle seed and gene databases were generated using reference sequences downloaded from NCBI (mitochondrial: NC_040987.1, ribosomal: AF169700.1). Mitochondrial assembles were visualised using Circos and a custom python script (https://github.com/o-william-white/circos_plot_organelle). To compare the utility of shotgun and targeted sequencing, quality filtered reads were mapped to the target sequences identified using BWA-mem. Mapped reads with a minimum mapping quality threshold of 30 were used to calculate the number of bases and number of reads which were on or off target using samtools bedcov (Li et al., 2009) and bedtools multcov (Quinlan & Hall, 2010) respectively. Target enrichment was estimated for each sample by calculating the ratio of bases and reads that were either on or off target, whilst accounting for differences in sample sequencing depth. Mapped reads from shotgun and exome capture with a minimum mapping and site quality of 30 were then used to call variants using bcftools (Danecek et al., 2021). Variants were further processed with vcftools (Danecek et al., 2011) to only include sites with the target sequences, with a minimum minor allele count of three, maximum number of alleles of two, and maximum of 50% missing data per variant. Mean depth and quality were calculated across sites using vcftools.

## Results

### PacBio IsoSeq sequencing

The number of high-quality isoforms generated using the SMRTlink pipeline were 115,591 and 248,047 for the male (29659_1) and female (29659_4) sample respectively. The number of low-quality isoforms was 18 and 40 for the male (29659_1) and female (29659_4) sample respectively. The transcript N50 ranged from 2,115 (29659_1) to 2,018 (29659_4). Each sample exhibited a similar proportions of BUSCO gene categories (Figure 2A). The majority of BUSCO genes identified as “complete and single copy” or “complete and duplicated” (61%) were unique to the female sample (29659_4; Figure 2B), with 37% shared and 4% unique to the male sample (29659_1; Figure 2B). The sequence taxonomy as defined by blobtools was most frequently identified as the class Malacostraca (Figure 2C).

**Figure 2.**
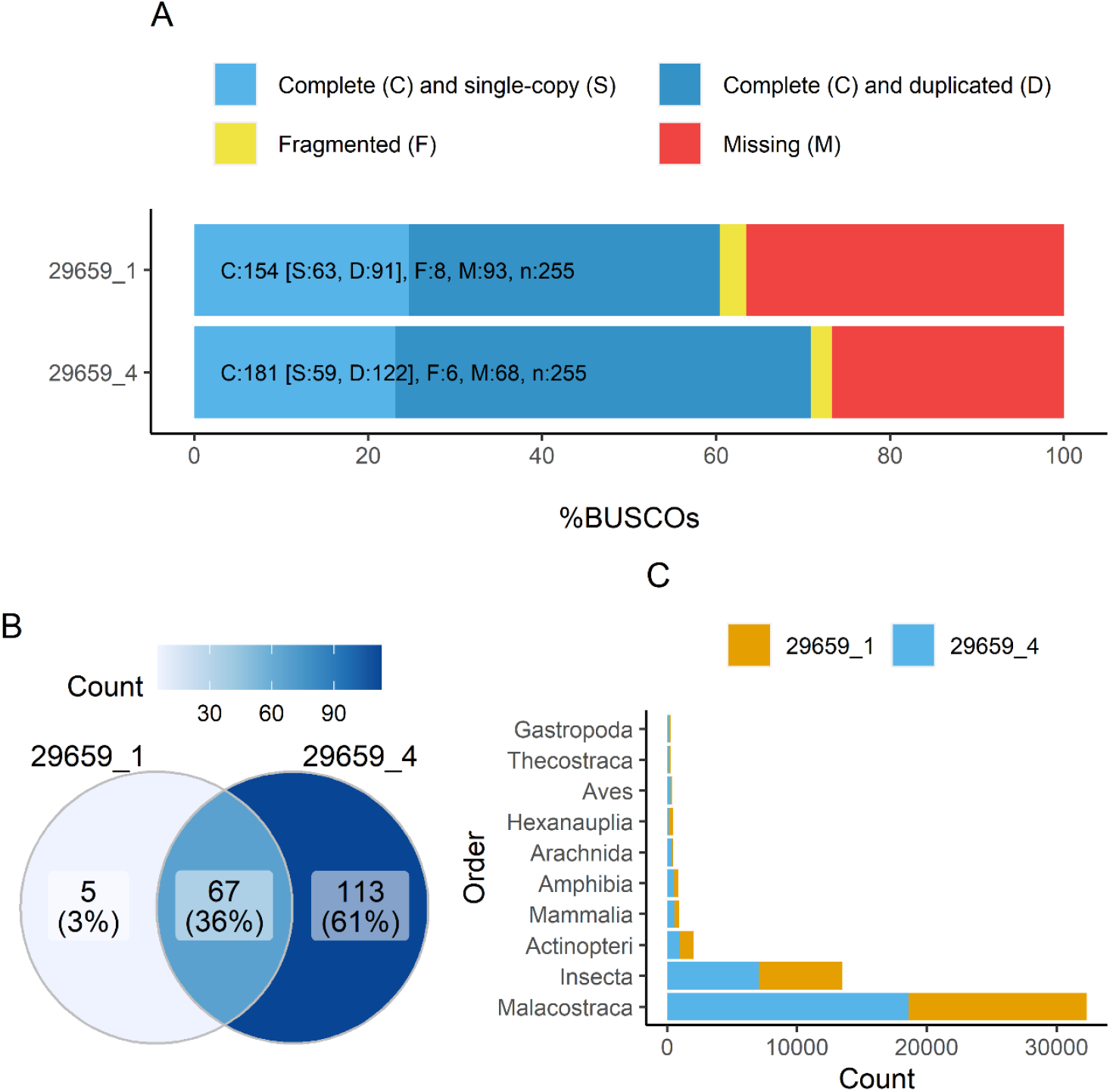
(A) Proportion of BUSCO categories identified, (B) Venn diagram of complete or duplicated BUSCO genes identified across samples and (C) number transcripts identified across the ten most frequent classses.

### Target selection and bait design

A total of 175 transcripts were initially selected as putative target sequences, identified as core BUSCO eukaryotic genes (64), having a blast description suggesting a role in environmental responses or core homeobox genes (65) or being identified as a gene from *Euphuasia superba* (46). Nine transcripts were removed for having a high similarity to other putative target sequences or being a putative repetitive sequence. A final set of 166 target sequences with a total target size of 395,924nt were selected and tested for bait deign suitability.

A total of 18,468 baits were designed using 80nt probes and 4× tiling. Of these, 8,244 baits passed filters based on blast searches against reference sequences and softmasking for repeats. A total of 43 target sequences were completely covered by baits, 120 targets were covered by baits with gaps up to 100nt and three targets had no baits. With this bait design, 301,711nt from a total of 395,924nt (∼75%) in the target sequences could be targeted.

### Shotgun and exome capture sequencing

A total of 46Mb and 17Mb of sequencing was generated for shotgun and exome capture sequencing respectively. Shotgun sequencing generated a mean of 15.4M raw reads and 5.5M quality filtered reads per sample. Exome capture generated a mean of 5.8M raw reads and 2.4M quality filtered reads per sample (Table 2). Of the 20 samples used for shotgun sequencing, GetOrganelle assembled mitochondrial sequences from 12 (60%) of samples, of which 6 were circular (30%; Figure 3). For the ribosomal sequences, sequences could be assembled for 10 (50%) of samples (Table 2).

**Table 2.** Summary for shotgun and target sequence data for raw data and quality-controlled reads, organelle and ribosomal assembled sequences and on and off target reads and bases.

| Ref. | Type | Raw reads | QC reads | Raw bases | QC bases | Mt. status | Mt. bases | Nr. Bases | On target reads | Off Target reads | On target bases | Off target bases |
| --- | --- | --- | --- | --- | --- | --- | --- | --- | --- | --- | --- | --- |
| K1 | shotgun | 18016900 | 2661036 | 2720551900 | 320540908 | circular | 16768 | 1288 | 7119 | 26587 | 137896 | 1044701 |
| K2 | shotgun | 20250526 | 8392788 | 3057829426 | 1236206937 | 6 contigs | 1917 | NA | 134 | 2525 | 1271 | 82617 |
| K3 | shotgun | 18254058 | 5509612 | 2756362758 | 796133516 | 3 contigs | 12970 | 828 | 1327 | 7790 | 20498 | 238739 |
| K4 | shotgun | 17599512 | 6273522 | 2657526312 | 920703883 | NA | NA | 306 | 865 | 4855 | 12064 | 138770 |
| K5 | shotgun | 33627108 | 1326736 | 5077693308 | 132519789 | NA | NA | NA | 504 | 58164 | 7627 | 1822480 |
| K6 | shotgun | 19055218 | 3152918 | 2877337918 | 401856419 | circular | 17936 | 1066 | 16448 | 62193 | 335985 | 2715620 |
| K7 | shotgun | 2965722 | 1819434 | 447824022 | 268706600 | NA | NA | NA | 0 | 144 | 0 | 4858 |
| K8 | shotgun | 10587052 | 1507242 | 1598644852 | 165253271 | circular | 17267 | 1048 | 4034 | 16480 | 82969 | 676405 |
| K9 | shotgun | 10142844 | 6814336 | 1531569444 | 1015195287 | NA | NA | NA | 13 | 824 | 463 | 27236 |
| K10 | shotgun | 12492450 | 6167292 | 1886359950 | 885148181 | 1 contig | 16182 | 766 | 3437 | 11843 | 70463 | 500749 |
| K11 | shotgun | 10598340 | 6853402 | 1600349340 | 1016422097 | NA | NA | NA | 1 | 231 | 14 | 7667 |
| K12 | shotgun | 19937660 | 4756106 | 3010586660 | 593914559 | circular | 17940 | 1186 | 4376 | 23827 | 87443 | 914699 |
| K13 | shotgun | 16524502 | 6489468 | 2495199802 | 950084592 | 9 contigs | 5429 | NA | 20 | 1366 | 215 | 46663 |
| K14 | shotgun | 19387356 | 12007924 | 2927490756 | 1754038099 | 1 contig | 16834 | 749 | 1474 | 4962 | 32717 | 249285 |
| K15 | shotgun | 13780532 | 5735282 | 2080860332 | 816844722 | circular | 16767 | 775 | 7695 | 23246 | 155776 | 1040720 |
| K16 | shotgun | 15048366 | 10189934 | 2272303266 | 1494990999 | 1 contig | 16065 | NA | 202 | 1233 | 5182 | 64500 |
| K17 | shotgun | 8299664 | 6046690 | 1253249264 | 897888181 | NA | NA | NA | 22 | 1126 | 463 | 38449 |
| K21 | shotgun | 13867176 | 8267784 | 2093943576 | 1223444992 | NA | NA | NA | 17 | 808 | 158 | 27222 |
| K24 | shotgun | 13687748 | 2822042 | 2066849948 | 354625476 | circular | 17949 | 1551 | 6936 | 21202 | 139830 | 938396 |
| K27 | shotgun | 13911002 | 3588248 | 2100561302 | 529495334 | NA | NA | NA | 15 | 1009 | 62 | 33194 |
| K1 | target | 8402058 | 1388300 | 1268710758 | 165009734 | - | - | - | 192129 | 96328 | 7238533 | 4446313 |
| K2 | target | 6696780 | 1903326 | 1011213780 | 271680266 | - | - | - | 8763 | 6538 | 170344 | 167938 |
| K3 | target | 8049242 | 1632944 | 1215435542 | 213381115 | - | - | - | 23710 | 16502 | 648904 | 476362 |
| K4 | target | 4227688 | 970570 | 638380888 | 137661749 | - | - | - | 8962 | 5060 | 191111 | 130223 |
| K5 | target | 6004434 | 384790 | 906669534 | 31606967 | - | - | - | 8023 | 24479 | 334267 | 756903 |
| K6 | target | 8942204 | 1371860 | 1350272804 | 166531526 | - | - | - | 282062 | 178888 | 11659601 | 8390632 |
| K7 | target | 6594460 | 5541342 | 995763460 | 822983663 | - | - | - | 2317 | 636 | 53365 | 21704 |
| K8 | target | 5597778 | 943988 | 845264478 | 120891220 | - | - | - | 215856 | 120775 | 8065079 | 5229059 |
| K9 | target | 3564986 | 2135396 | 538312886 | 318452554 | - | - | - | 5065 | 591 | 205324 | 19536 |
| K10 | target | 4996168 | 2732408 | 754421368 | 399627439 | - | - | - | 183486 | 81742 | 7003341 | 3606622 |
| K11 | target | 7644834 | 6699940 | 1154369934 | 998112478 | - | - | - | 6213 | 1961 | 120675 | 76989 |
| K12 | target | 5141130 | 1056672 | 776310630 | 120100423 | - | - | - | 56073 | 30550 | 1919386 | 1254959 |
| K13 | target | 3866136 | 1504502 | 583786536 | 218348768 | - | - | - | 8398 | 2128 | 192086 | 53202 |
| K14 | target | 9276902 | 5932822 | 1400812202 | 876297636 | - | - | - | 310157 | 163433 | 11358129 | 7700727 |
| K15 | target | 5048616 | 2153850 | 762341016 | 303094169 | - | - | - | 204996 | 108658 | 8940070 | 5562644 |
| K16 | target | 7829856 | 6280716 | 1182308256 | 933550565 | - | - | - | 142909 | 64512 | 4494206 | 3185313 |
| K17 | target | 2248172 | 1592734 | 339473972 | 236913855 | - | - | - | 9655 | 2341 | 211240 | 86333 |
| K21 | target | 2970194 | 2359252 | 448499294 | 350268321 | - | - | - | 7971 | 2441 | 135877 | 65219 |
| K24 | target | 6203096 | 2277690 | 936667496 | 319810881 | - | - | - | 552622 | 315401 | 23460662 | 16415042 |
| K27 | target | 2876392 | 477518 | 434335192 | 68606137 | - | - | - | 995 | 1100 | 16702 | 30287 |

**Figure 3.**
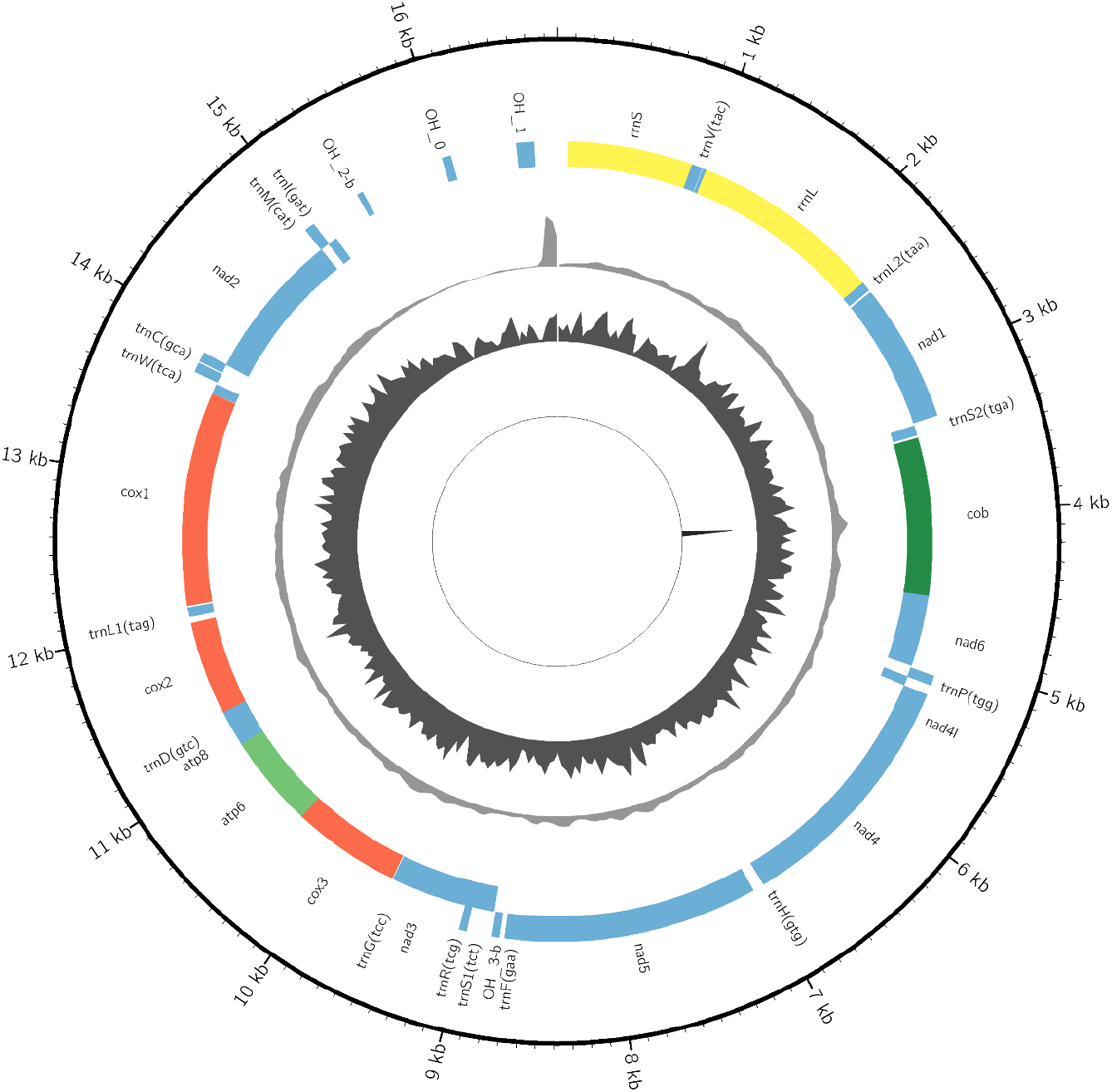
Circular mitochondrial sequence for sample K1 (9473-JK-1_001) visualised using circus and showing the following attributes from outside to inside: sequence position, annotation names, annotations on the + strand, annotations on the - strand, coverage, GC content, repeat content.

The mean ratio of on target:off target bases was 0.07 and 2.07 for shotgun and exome capture sequencing respectively, representing mean fold change of ∼30× for targeted sequencing (Figure 4A). The mean ratio of on target:off target reads was 0.14 and 2.36 for shotgun and exome capture sequencing respectively, resulting in a mean fold change of ∼16× (Figure 4B). Following variant calling and filtering, a total of 28 and 75 variants were identified for the shotgun and target sequence data. The variant mean depth was significantly higher for targeted sequencing when compared to shotgun sequencing. In addition, site quality was higher for targeted sequencing.

**Figure 4.**
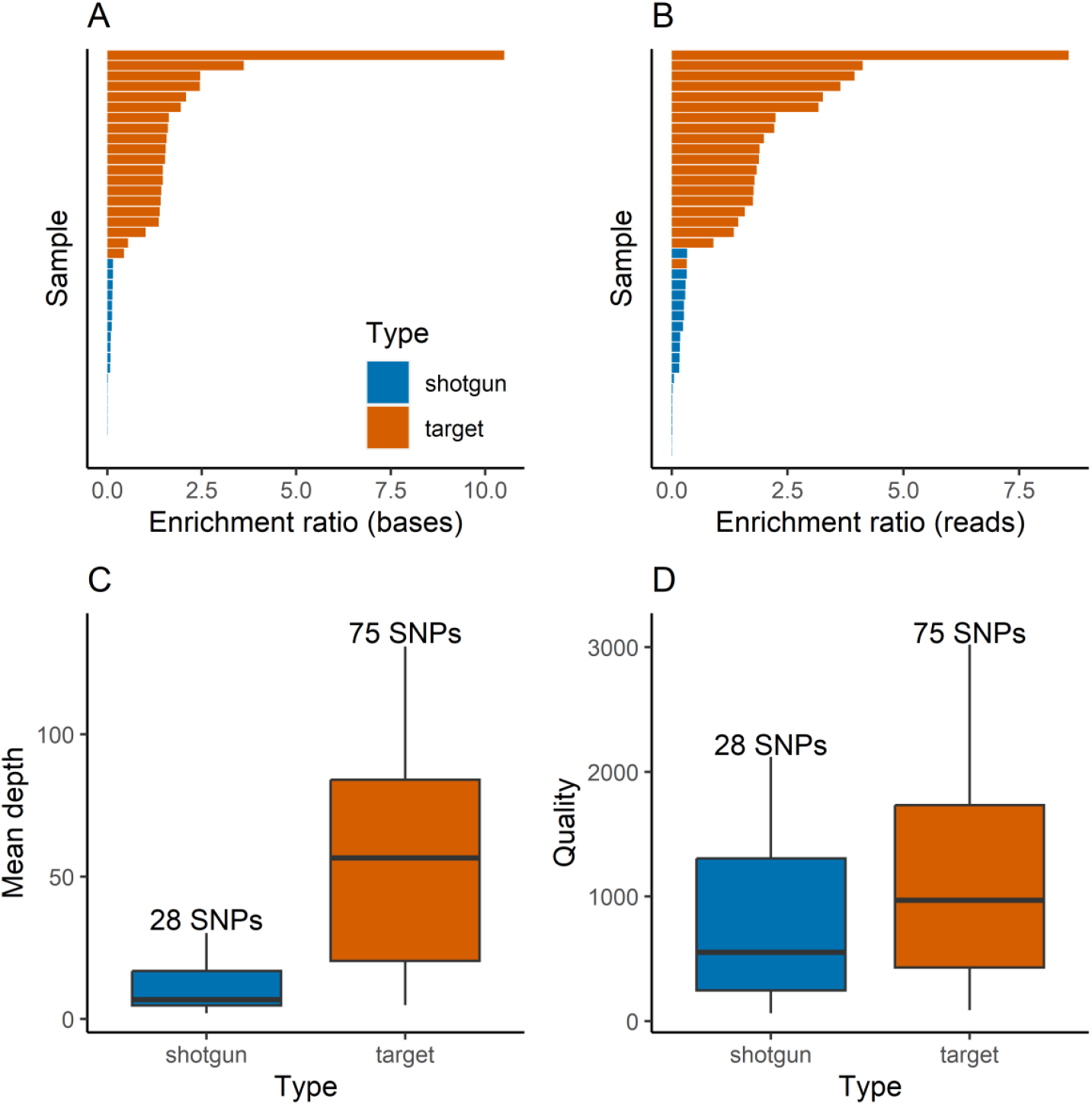
Enrichment of mapped (A) bases (B) reads calculated a ratio of on:off target mapping. (C) Mean depth and (D) quality of called sites. Samples in plots A and B are sorted by enrichment ratio.

## Discussion

This study assessed the relative utility of shotgun and exome capture sequencing for genomic analyses with historical spirit collections of Antarctic krill. Importantly, this study highlights that it is possible to isolate DNA from historical krill spirit collections and that these samples can be used for genomic analyses. This study has important implications for the utility of historical spirit collections, including the Discovery collections of the 1920s and 30s, unlocking the collections for genomic studies that could investigate changes in diversity and resilience of krill. For example, future work could investigate (1) what historical krill diversity looked like prior to the onset of anthropogenic climate change and widespread fishing, (2) how contemporary krill diversity been impacted by changes in climate change and predation pressure, (3) the relationship between the biological characteristics of krill and temperature and (4) how adaptable krill are to climatic change. Considering the central importance of krill in the food web of Southern, the implications for conservation and fisheries management. are potentially profound. Although, our study has focused on Antarctic krill, the approach could also be applied to a broader range of taxa.

Full length mitochondrial sequences and ribosomal sequences could be assembled using shotgun sequence data from historical samples. Multicopy parts of the genome such as the multicopy organelle genomes or ribosomal tandem repeats are common targets for “genome skimming” studies, as these regions are sequenced at a higher depth compared to the rest of the genome (Straub et al., 2012). This approach is useful to confirm that we have endogenous sequence data from the target species. Although variation in mitochondrial and ribosomal genes was identified, it is unlikely that these regions will have sufficient polymorphism required the resolve differences, if any, in krill population structure.

Although our study highlights that exome capture can be successfully applied to historical spirit collections, there are clear differences in DNA sample and sequence quality. Spirit or wet collections are typically stored in liquid preservatives including ethanol and may have been fixed with formalin prior to storage which may damage DNA (Ruiz-Gartzia et al., 2022). Recent studies investigating the utility of spirit collections for genomic studies have highlighted that overall specimen condition had the greatest impact on recovering high quality genomic DNA (Hahn et al., 2022; O’Connell et al., 2022; Straube et al., 2021).

Concerning the selection and design of bait probes, our analysis highlighted potential issues for future studies. Given the repetitive nature of the krill genome, we will be unable to design baits for some regions because of non-specific binding. Specifically, we were able to design baits for 76% of nucleotides in the original gene target list. Therefore, future studies must account for the likelihood of bait deign failure for target sequences. In addition, our study highlighted that sequence depth was often higher at the boundaries of target sequences where bait frequency was lower due to the existence of repetitive sequences. To reduce the likelihood of sequencing repetitive sequences, buffer zones of around repetitive sequences could be identified and avoid designing baits for these regions. In additions, baits could be designed for regions with a minimum depth of bait sequence (for example 3×).

## Acknowledgements

The authors would like to acknowledge Gavin Horsburgh, Lucy Knowles (The University of Sheffield) and Xuan Liu (The University of Liverpool) for the sample preparation, RNA extraction and bioinformatic processing for the PacBio RNAseq data. The authors would also like to acknowledge Jennifer Klunk from Arbor biosciences for technical support and advice regarding Arbor biosciences services. We would also like to acknowledge project collaborators Melody Clark, Simeon Hill (British Antarctic Survey) and Thomas Mock (University of East Anglia) for comments on the manuscript.

## Author contributions

OW and MC designed the study. GT provided the samples required for full length transcriptomes and LH provided the historical samples for shotgun and exome capture sequencing. SW performed the wet lab work. OW wrote the first draft of the manuscript; all co-authors contributed to the preparation of the final manuscript.

